# Distinct Roles of ADP and Magnesium in the Structural Remodeling and Higher-Order Assembly of Peroxiredoxin 2

**DOI:** 10.64898/2026.11.07.765024

**Authors:** Tran Ngoc Trang, Ryusei Yamada, Koki Morita, Hiroki Konno

**Author notes:** Correspondence and request for materials should be addressed to HK. These authors contributed equally to this work (Tran Ngoc Trang, Ryusei Yamada).

## Abstract

2-Cys peroxiredoxins (Prxs) undergo oligomeric transitions linked to peroxidase and chaperone activities, but the contributions of nucleotides and cations to lipid-dependent assembly remain unclear. We examined phosphatidylserine (PS)-induced Prx2 oligomers using biochemical assays and high-speed atomic force microscopy (HS-AFM). Both wild-type (WT) and P44A/S76A mutant (MT) Prx2 formed chaperone-active oligomers, although MT assembled more slowly and exhibited a more compact lateral geometry. Mg^2+^ disassembled both oligomers, whereas ADP alone predominantly disassembled WT. Under combined Mg-ADP conditions, HS-AFM directly captured oligomer disassembly in both proteins, and separate experiments demonstrated higher-molecular-weight (HMW) assembly. WT HMW assemblies exhibited greater heights and stronger Mant-ADP co-elution than MT. Structural predictions suggested altered local Ser76-mediated contacts without major backbone changes. These results distinguish PS-induced functional switching from Mg^2+^- and ADP-dependent remodeling and support, without directly proving, a disassembly–reassembly route to HMW formation.

## INTRODUCTION

Peroxiredoxins (Prxs) are conserved thiol-dependent peroxidases that protect cells from oxidative stress. Mammalian Prx2 is a major cellular antioxidant [1-3]. Beyond peroxidase activity, 2-Cys Prxs switch among dimers, decamers, and larger assemblies, sometimes acquiring molecular chaperone activity [2-7]. Chaperone-active Prxs support stress tolerance and proteostasis in yeast and *Leishmania* [4, 8-10], although these assemblies need not share a common structure or function. We previously showed that anionic phospholipids, including phosphatidylserine (PS), convert human Prx2 into trefoil-shaped, chaperone-active oligomers with diminished peroxidase activity [11, 12]. Mg-ADP subsequently promotes spherical high-molecular-weight (HMW) assemblies [11]. PS also influences Prx2 localization and chaperone function on exosomes [13], but cellular formation of these spherical HMW structures has not been established. How Mg^2+^ and ADP individually affect PS-induced oligomers, and whether HMW formation involves remodeling of existing oligomers, remain unknown. We compared WT Prx2 with a P44A/S76A mutant (MT) using biochemical assays and high-speed atomic force microscopy (HS-AFM) to distinguish the effects of Mg^2+^ and ADP on lipid-dependent oligomer remodeling and HMW assembly.

## RESULTS AND DISCUSSION

### P44A/S76A substitutions reduce nucleotide association with Prx2 HMW assemblies

To examine how nucleotide-related interactions influence Prx2 assembly, we first used ATPbind to identify candidate nucleotide-interacting residues. Pro44 and Ser76 showed relatively high prediction scores (Fig. 1A), and we prepared the P44A/S76A double mutant (MT) for comparison with wild-type (WT) Prx2 (Fig. S1). These residues were selected computationally; their direct involvement in nucleotide binding has not been established. We analyzed assembly by size-exclusion chromatography (SEC), which separates species according to hydrodynamic size. Neither phosphatidylserine (PS) with Mg^2+^/ADP/Mant-ADP nor WT Prx2 with Mg^2+^/ADP/Mant-ADP produced an early-eluting high-molecular-weight (HMW) peak. In contrast, a pronounced peak appeared when Prx2, PS, and Mg^2+^/ADP/Mant-ADP were combined (Fig. 1B). Thus, these components are jointly required under the conditions tested, consistent with our previous report [11]. Under HMW-forming conditions, MT also generated an HMW fraction, but the peak was less pronounced than that of WT (Fig. S2). Mant-ADP fluorescence strongly co-eluted with the WT HMW fraction and much more weakly with the MT fraction (Fig. 1C). The reduced HMW peak and weaker Mant-ADP co-elution observed for MT suggest effects of the P44A/S76A substitutions on both HMW assembly and nucleotide association. However, co-elution alone does not identify a direct binding site or establish binding stoichiometry.

**Fig. 1.**
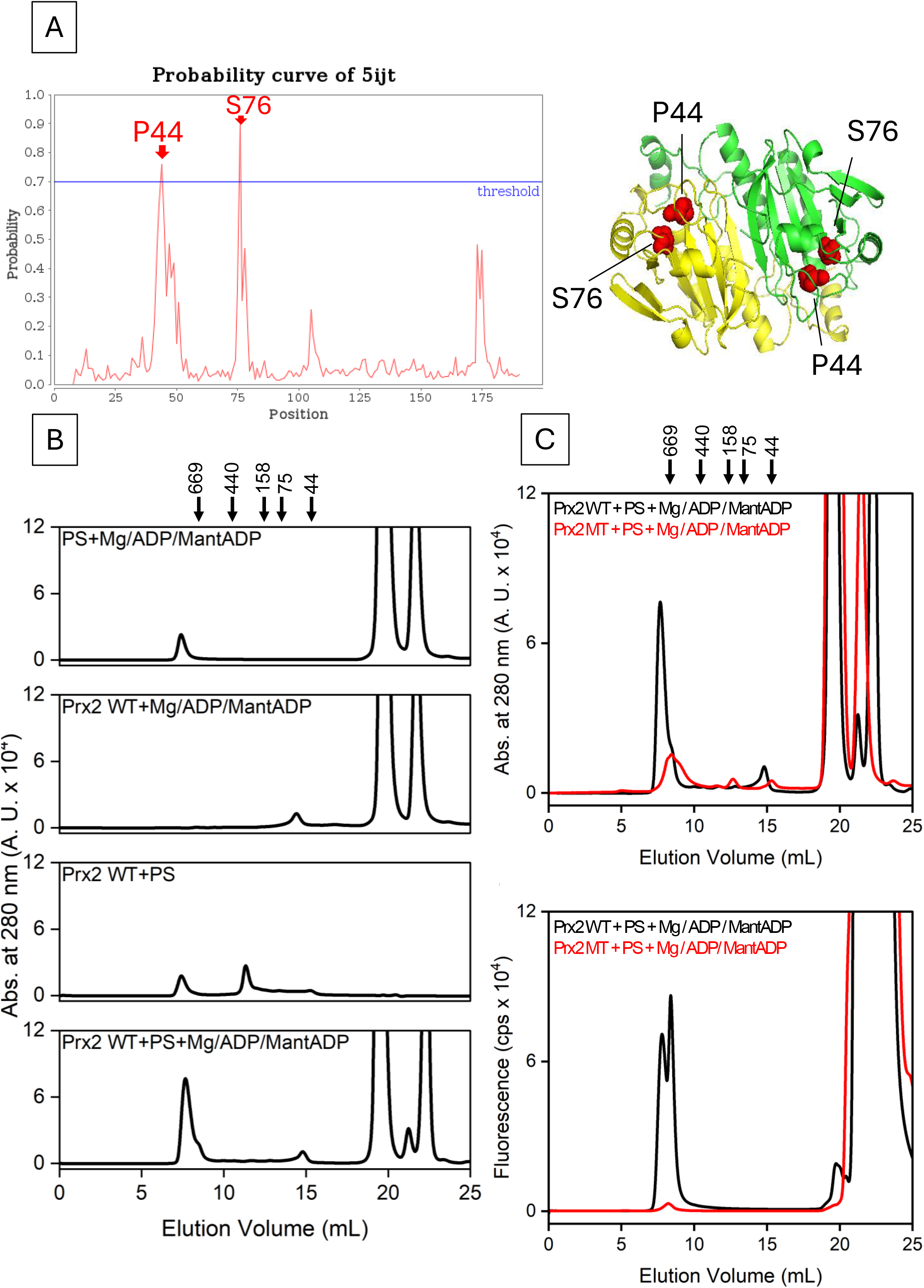
Effects of P44A/S76A substitutions on Mg-ADP-dependent HMW assembly of Prx2. (A) ATPbind prediction of nucleotide-interacting Prx2 residues (threshold 0.7; https://zhanggroup.org/ATPbind/). P44 and S76 are highlighted on PDB 5ijt. (B) SEC profiles of PS plus Mg^2+^/ADP/Mant-ADP and WT Prx2 with Mg^2+^/ADP/Mant-ADP, PS, or both. An early-eluting HMW peak appears only with Prx2, PS, and nucleotide-containing mixture; absorbance at 280 nm. (C) WT (black) and MT (red) SEC absorbance (280 nm, top) and Mant-ADP fluorescence (355/448 nm, bottom) under HMW-forming conditions. MT shows weaker HMW-associated fluorescence.

### P44A/S76A substitutions alter the architecture and assembly behavior of PS-induced Prx2 oligomers

We next asked whether the mutant already differed from WT before exposure to Mg^2+^ and ADP. Following incubation with PS, both proteins produced SEC peaks at 11-13 mL, confirming that MT retains PS-dependent oligomerization (Fig. 2A and 2B). High-speed atomic force microscopy (HS-AFM) showed trefoil-shaped oligomers for both proteins (Fig. 2C; Movies S1 and S2). Nevertheless, the triangular area measured across the three lobes was smaller for MT than WT (11.97 ± 4.41 versus 14.18 ± 5.78 nm^2^); MT oligomers also showed a slightly greater height (Fig. S3). These differences indicate a change in the observed oligomer geometry, rather than loss of oligomer formation. Time-course SEC further revealed that WT oligomerization approached a plateau within approximately 1 h, whereas MT assembled more slowly and reached a lower oligomer yield (Fig. 2D). Thus, the mutations affect both the architecture and formation efficiency of the PS-induced state. Importantly, the persistence of a trefoil-shaped morphology in MT indicates that P44A/S76A does not prevent the basic PS-driven assembly pathway but modifies the geometry and population of the resulting oligomers.

**Fig. 2.**
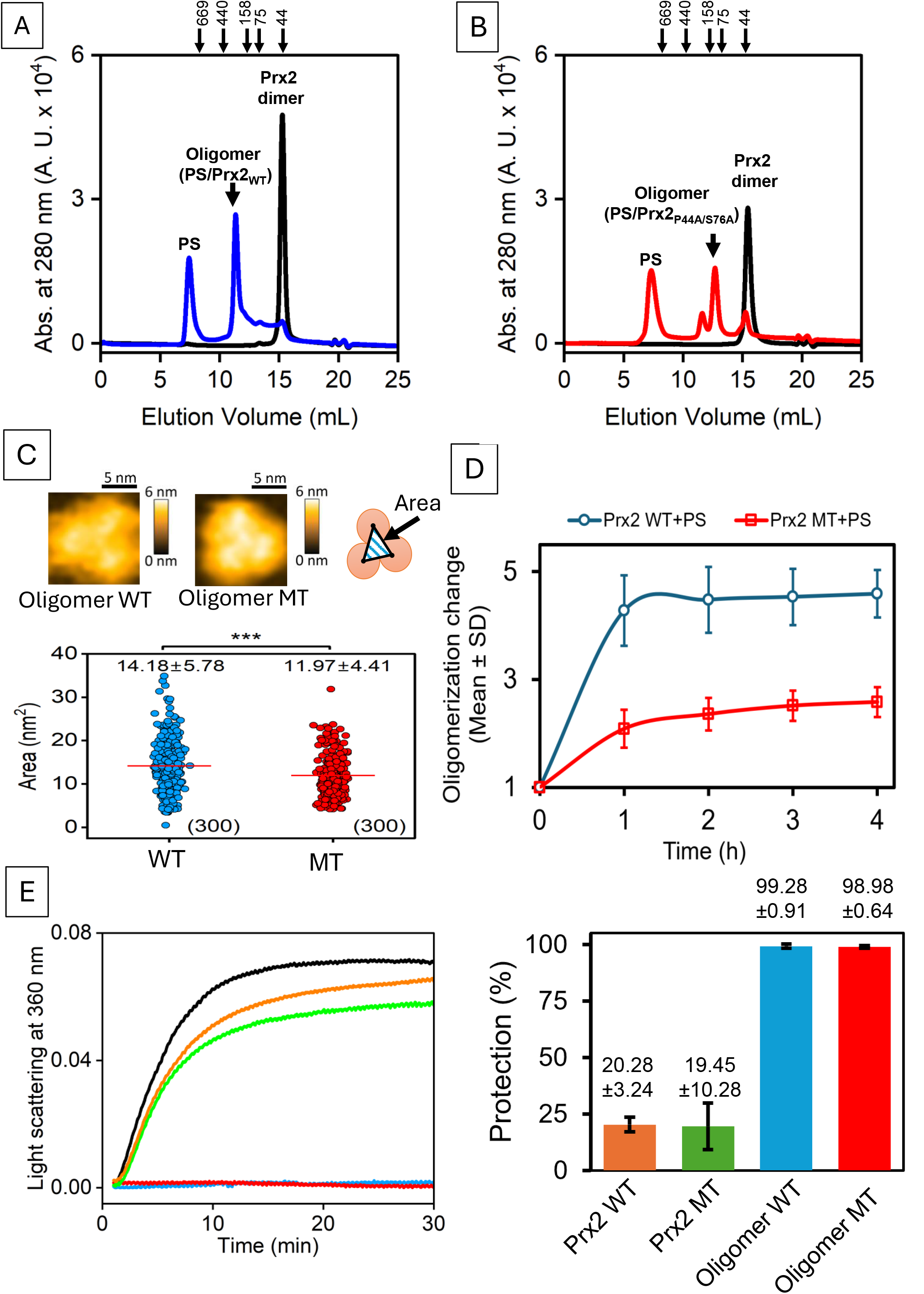
Structural and functional comparison of Prx2 WT and Prx2 MT oligomers. (A,B) SEC profiles of WT (A) and MT (B) after 2 h with PS (blue/red) or without PS (black). (C) HS-AFM analysis of trefoil oligomer geometry. Triangular vertices correspond to the highest points of the three lobes. WT (blue) and MT (red) area distributions; red horizontal lines indicate means. ***p < 0.001, Student’s t-test. (D) SEC time course of PS-induced oligomerization. WT plateaus near 1 h; MT accumulates more slowly and to a lower level. Mean ± SD. (E) Citrate synthase (CS) thermal aggregation monitored by scattering at 360 nm: CS alone (black), non-oligomeric WT (orange), MT (green), oligomeric WT (blue), and MT (red). Both oligomers provide approximately 99% protection at the tested concentration.

We then assessed whether these differences compromised chaperone function using the thermal aggregation of citrate synthase (Fig. S4). PS-induced WT and MT oligomers both strongly suppressed aggregation, whereas non-oligomeric samples provided substantially weaker protection (Fig. 2E). No significant difference between WT and MT HMW chaperone activities was detected under the tested conditions (Fig. S5). Therefore, changes in oligomer geometry and assembly efficiency do not necessarily abolish PS-dependent chaperone activation; the assay does not exclude differences at other protein concentrations. The separation between altered assembly behavior and largely preserved protection against citrate synthase aggregation suggests that the structural requirements for oligomer formation and chaperone activity are not identical. To explore a possible molecular basis, we superimposed predicted WT and MT structures (Table 1; Figs. S6 and S7). The predicted backbones were nearly identical (RMSD 0.07 Å), including near residues 44 and 76. In the WT model, the Ser76 hydroxyl lies near Tyr43, Asp46, and Asp78 at distances compatible with potential hydrogen bonds. Substitution with alanine removes this hydroxyl and its predicted local contacts without a major modeled backbone rearrangement. These structural predictions may explain the altered oligomer organization and ADP responsiveness, although the PS-induced oligomer interface and the individual contributions of the two substitutions remain unresolved.

### Mg^2+^ and ADP induce distinct disassembly responses in PS-induced Prx2 oligomers

We next examined how preformed PS-induced oligomers respond to Mg^2+^ and ADP separately (experimental sequence in Fig. S8). Mg^2+^ reduced the oligomer SEC peaks of both WT and MT and increased later-eluting smaller species (Fig. 3A). Time-lapse HS-AFM directly visualized fragmentation of both trefoil-shaped oligomers (Fig. 3B; Movies S3 and S4). Thus, Mg^2+^ can destabilize both oligomer architectures. Agreement between SEC and direct HS-AFM imaging is important here: the chromatographic shift indicates a change in the distribution of assembly states, while imaging shows that individual preformed oligomers actually break apart. ADP alone produced a different pattern: WT oligomer peaks decreased and smaller species appeared, whereas MT oligomers remained largely unchanged (Fig. 3C). HS-AFM independently showed disassembly of WT but substantially greater persistence of MT oligomers (Fig. 3D; Movies S5 and S6). The contrasting responses to ADP and Mg^2+^ suggest that P44A/S76A selectively alters ADP-dependent structural remodeling rather than generally stabilizing Prx2 oligomers against disassembly. Because the mutant still responds to Mg^2+^, the two components cannot be assumed to act through an identical destabilization mechanism.

**Fig. 3.**
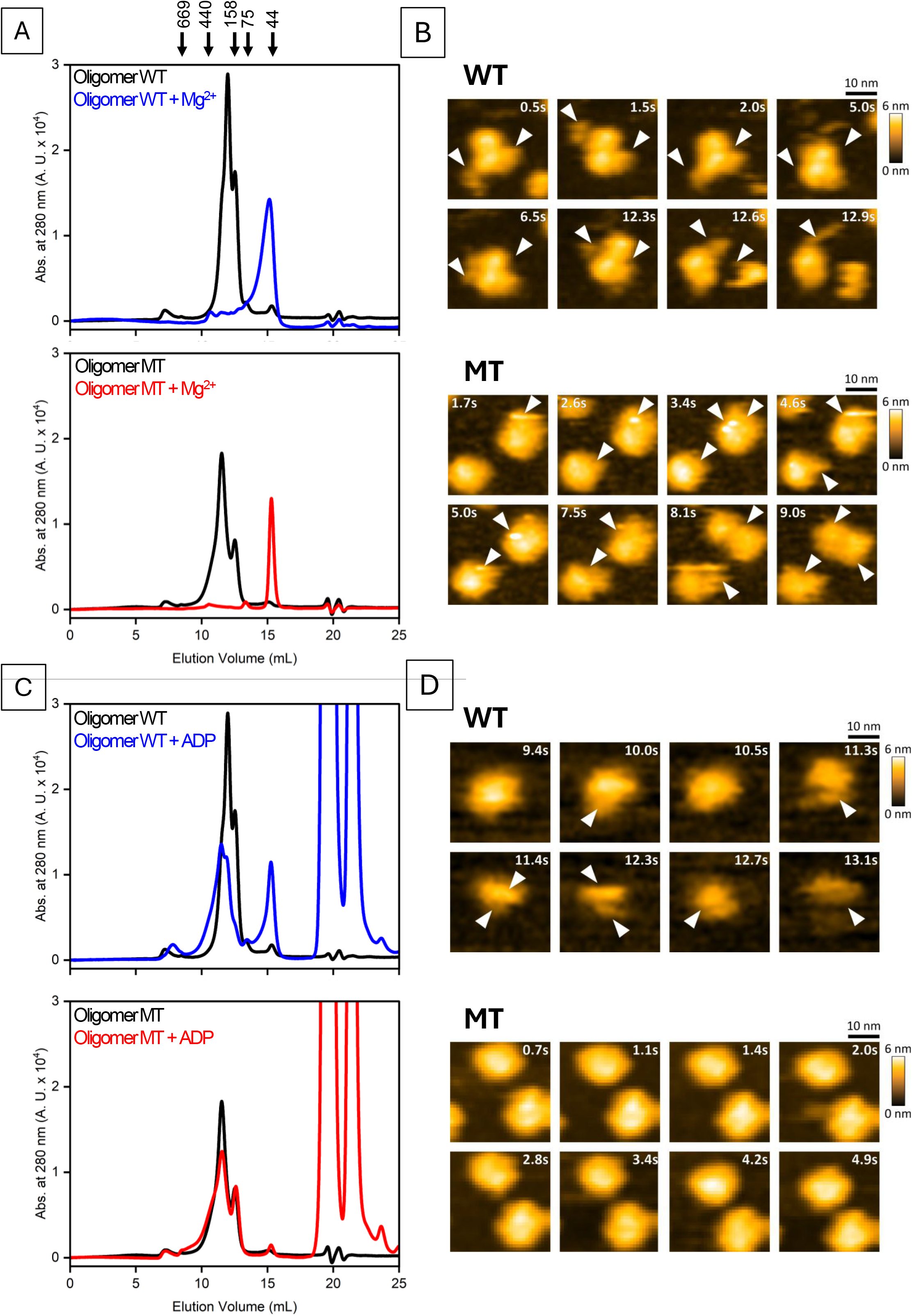
Mg^2+^ and ADP trigger distinct disassembly pathways of Prx2 oligomers. (A) SEC profiles before (black) and after 12 h with MgCl_2_ (WT blue, MT red), showing later-eluting smaller species. (B) HS-AFM movies of Mg^2+^-induced oligomer disassembly. Arrowheads mark fragmentation. Scale bar 10 nm; Z-scale 6 nm; scan 50 × 50 nm^2^, 50 × 50 pixels. (C) SEC profiles after 12 h with ADP. WT oligomers partly disassemble; MT remains largely unchanged. (D) HS-AFM imaging under ADP. Arrowheads mark fragmentation. Scale bar 10 nm; Z-scale 6 nm; scan 50 × 50 nm^2^, 50 × 50 pixels.

Importantly, when Mg^2+^ and ADP were present together, HS-AFM directly captured disassembly of pre-existing PS-induced oligomers in both WT and MT (Fig. S9; Movies S7 and S8). This observation demonstrates that disassembly also occurs under the combined condition used for HMW formation. The distinct responses to ADP and Mg^2+^ suggest that ADP sensitivity depends on structural features affected by P44A/S76A, while Mg^2+^ promotes destabilization of both oligomer types. The reduced response of MT to ADP does not exclude nucleotide binding, because oligomer disassembly reflects both nucleotide association and the subsequent structural rearrangements. Consistent with this interpretation, Mant-ADP co-eluted less efficiently with MT HMW assemblies.

### WT and MT exhibit distinct HMW assembly under Mg-ADP conditions

Finally, we characterized the HMW assemblies obtained in the presence of Mg^2+^ and ADP. SEC detected early-eluting HMW fractions for both WT and MT, showing that neither P44 nor S76 is strictly required for assembly under these conditions (Fig. 4A). Mant-ADP fluorescence co-eluted strongly with WT HMW and weakly with MT HMW (Fig. 4B), consistent with reduced nucleotide association in MT. HS-AFM showed that both assemblies developed vertically over time, but WT reached greater heights: 27.0 ± 12.4 versus 15.4 ± 5.3 nm at 4 h, and 51.0 ± 27.4 versus 35.0 ± 14.6 nm at 16 h for WT and MT, respectively (Fig. 4C and 4D). These measurements describe AFM height, not molecular mass or total particle volume. They indicate that the substitutions affect the development of HMW architecture as well as nucleotide co-elution. Our findings support the model in Fig. 5: PS first promotes chaperone-active oligomers; Mg^2+^ and ADP then induce structural remodeling, including disassembly, under conditions that also produce spherical HMW assemblies.

**Fig. 4.**
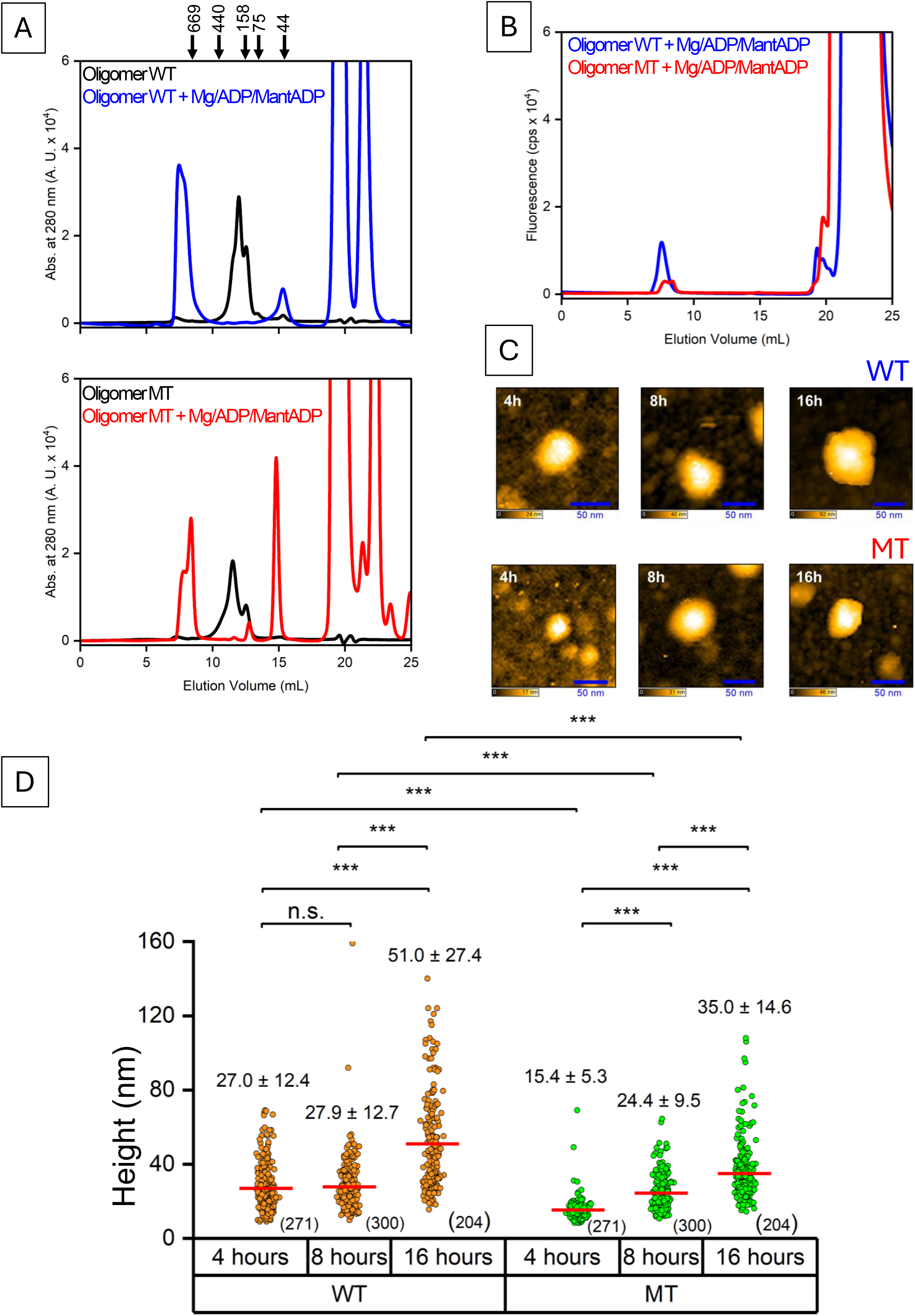
Distinct HMW assembly properties and Mant-ADP association of WT and MT Prx2 under Mg-ADP conditions. (A) SEC absorbance profiles of WT (top) and MT (bottom) HMW assemblies after 12 h with Mg^2+^/ADP/Mant-ADP. (B) Mant-ADP fluorescence co-elution with WT (blue) and MT (red) HMW fractions. (C) HS-AFM images of HMW assemblies at indicated times; scale bar 50 nm, scan 300 × 300 nm^2^. (D) Height distributions of WT (orange) and MT (green) HMW assemblies.

**Fig. 5.**
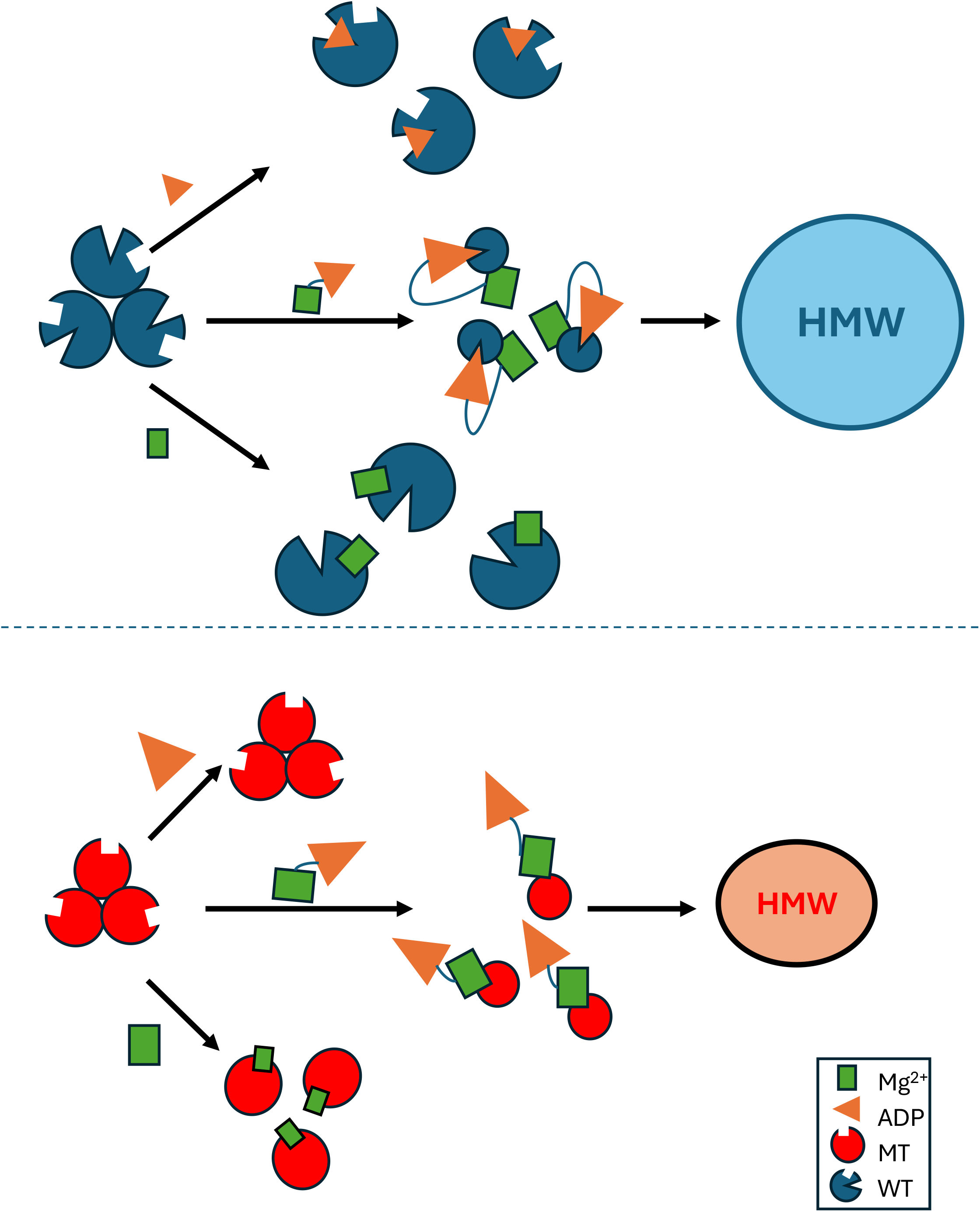
Proposed model of Mg^2+^- and ADP-dependent structural remodeling and HMW assembly of Prx2. Proposed structural remodeling of PS-induced WT (blue) and MT (red) oligomers. Mg^2+^ (green squares) disassembles both, whereas ADP (orange triangles) predominantly disassembles WT. Combined Mg-ADP promotes spherical HMW assembly in both, with lower heights and weaker Mant-ADP co-elution in MT. The disassembly–reassembly route is proposed, not directly tracked in individual molecules.

HS-AFM directly captured Mg-ADP-induced disassembly, whereas HMW formation was measured separately, and the same dissociated molecules were not followed continuously into HMW structures. The proposed disassembly– reassembly sequence is therefore a model, not an established single-molecule trajectory. The P44A/S76A substitutions caused little predicted backbone rearrangement but altered local Ser76-mediated hydrogen bond contacts (Table 1; Figs. S6 and S7), suggesting that local interactions may influence how PS-induced oligomers respond to ADP.

The physiological role of these PS- and nucleotide-dependent spherical HMW assemblies remains unresolved. Chaperone-active states of other Prxs contribute to proteostasis and stress tolerance [4, 8-10], although their oligomeric structures are not necessarily equivalent to those studied here. PS binding also regulates Prx2 localization and chaperone function on exosomes [13], providing a biological context for lipid-dependent regulation. Whether spherical HMW assemblies form in cells, and how physiological nucleotide concentrations or the order of component addition affect them, require further investigation.

## CONCLUSIONS

We show that PS-induced Prx2 oligomers undergo distinct responses to Mg^2+^ and ADP: Mg^2+^ destabilizes both WT and P44A/S76A oligomers, whereas ADP alone predominantly destabilizes WT. Under combined Mg-ADP conditions, HS-AFM directly reveals disassembly of both oligomer types, and both generate HMW assemblies with different heights and Mant-ADP co-elution. Predicted local interactions around Ser76 may contribute to these differences without substantial backbone rearrangement. These results identify structural remodeling as an important step in lipid-dependent Prx2 higher-order assembly, while the proposed reassembly pathway and physiological role of spherical HMW complexes remain to be established.

Both proteins formed HMW assemblies under Mg-ADP conditions, showing that P44 and S76 are not strictly required. Nevertheless, WT showed stronger Mant-ADP co-elution and greater HMW height (Fig. 4), suggesting that precursor oligomer architecture influences subsequent higher-order assembly (Fig. 5). Co-elution does not establish direct nucleotide-binding stoichiometry or site.

Our results distinguish PS-induced chaperone activation from subsequent Mg^2+^- and ADP-dependent structural remodeling. Direct imaging of Mg-ADP-induced disassembly, together with separate HMW assembly measurements, supports a disassembly–reassembly model without establishing a continuous molecular trajectory.

## Supporting information

Movie S1_Time-lapse HS-AFM movie showing structural dynamic of oligomer WT

Movie S2_Time-lapse HS-AFM movie showing structural dynamic of oligomer MT

Movie S3_Time-lapse HS-AFM movie showing the degradation of WT oligomers in the presence of MgCl2

Movie S5_Time-lapse HS-AFM movie showing the degradation of WT oligomers in the presence of ADP

Movie S6_Time-lapse HS-AFM movie of the interaction between MT oligomers and ADP

Movie S7_Time-lapse HS-AFM movie showing the degradation of WT oligomers in the presence of Mg-ADP

Movie S8_Time-lapse HS-AFM movie showing the degradation of MT oligomers in the presence of Mg-ADP

## Acknowledgments

We thank Prof. T. Ando of Kanazawa University for the critical reading of the manuscript. This work was supported by a KAKENHI grant for Research on Innovative Area (#25112507 to H.K.), WISE Program for Nano-Precision Medicine, Science, and Technology of Kanazawa University by MEXT, the World Premier International Research Center Initiative (WPI), MEXT, Japan, and the Kanazawa University CHOZEN project.

## Author contributions

H.K., T.N.T, and R.Y. designed the research; T.N.T, R.Y., and K.M. performed the biological experiments; T.N.T, and R.Y. performed the HS-AFM observations; T.N.T, and R.Y. analyzed the data. H.K., T.N.T. wrote the paper with input from all authors.

## The authors declare no conflict of interest

## Notes

### Competing Interest Statement

The authors have declared no competing interest.

