## Supporting information for "Distinct Roles of ADP and Magnesium in the Structural Remodeling and Higher-Order Assembly of Peroxiredoxin 2"

**This PDF file includes**


| | | |
|---|---|---|
| | Materials and Methods | Page S-3-4 |
| Figure S1 | Sequence alignment between Prx2 WT and Prx2 MT. | Page S-5 |
| Figure S2 | SEC chromatograms of the Prx2 MT under different conditions. | Page S-6 |
| Figure S3 | Height analysis of Prx2 oligomers. | Page S-7 |
| Figure S4 | Scheme illustrating the principle of the citrate synthase aggregation assay. | Page S-8 |
| Figure S5 | Chaperone activities of WT and MT Prx2 HMW assemblies. | Page S-9 |
| Table 1 | Data sets of structural comparison of AlphaFold-predicted WT and P44A/S76A hPrx2. | Page S-10 |
| Figure S6 | Structural comparison of AlphaFold-predicted WT and P44A/S76A hPrx2. | Page S-11 |
| Figure S7 | Backbone bending angle comparison of (A) Prx2 WT and (B) Prx2 MT. | Page S-12 |
| Figure S8 | Experimental procedure for confirming ADP and $Mg^{2+}$ involvement in HMW complex formation. | Page S-13 |
| Figure S9 | HS-AFM time-lapse imaging of oligomers in the presence of Mg-ADP, capturing progressive structural dissociate over time. | Page S-14 |
| Movie S1 | Time-lapse HS-AFM movie showing structural dynamic of oligomer WT. | Page S-15 |
| Movie S2 | Time-lapse HS-AFM movie showing structural dynamic of oligomer MT. | Page S-15 |
| Movie S3 | Time-lapse HS-AFM movie showing the degradation of WT oligomers in the presence of $MgCl_2$. | Page S-15 |
| Movie S4 | Time-lapse HS-AFM movie showing the degradation of MT oligomers in the presence of $MgCl_2$. | Page S-15 |

| | | |
|---|---|---|
| Movie S5 | Time-lapse HS-AFM movie showing the degradation of WT oligomers in the presence of ADP. | Page S-15 |
| Movie S6 | Time-lapse HS-AFM movie of the interaction between MT oligomers and ADP. | Page S-15 |
| Movie S7 | Time-lapse HS-AFM movie showing the degradation of WT oligomers in the presence of Mg-ADP. | Page S-15 |
| Movie S8 | Time-lapse HS-AFM movie showing the degradation of MT oligomers in the presence of Mg-ADP. | Page S-15 |

**Materials & Methods**

*Cloning, expression, and purification of the wild-type Prx2 and Prx2P$_{44A/S76A}$ mutant.*

The pET23a plasmid encoding human peroxiredoxin 2 was prepared as described previously [R1]. The P44A/S76A mutant, in which Pro44 and Ser76 were substituted with Ala, was generated using the QuikChange Site-Directed Mutagenesis method. Prx2 wild-type (WT) and the Prx2 P44A/S76A mutant (MT) were expressed in E. coli BL21 (DE3) using a previously described Prx2 expression strategy [R1]. For WT purification, cell pellets were resuspended in 50 mM sodium phosphate buffer (pH 7.2), 1 mM EDTA, and 40% ammonium sulfate, disrupted using a French press, clarified by centrifugation (15,000 rpm, 10 min, 4oC), and applied to a Toyopearl Phenyl hydrophobic interaction column (Tosoh, Tokyo, Japan). WT protein was eluted with a 20-0% ammonium sulfate gradient. MT samples were processed following the same overall procedure but without ammonium sulfate in the initial resuspension buffer. After disruption and clarification, the lysate was first subjected to Toyopearl DEAE anion exchange chromatography using a 0-400 mM NaCl gradient, then adjusted to 20% ammonium sulfate and purified by Toyopearl Butyl-650M hydrophobic interaction chromatography, from which MT protein was eluted with a 20-0% ammonium sulfate gradient. Both WT and MT proteins were subsequently purified by size-exclusion chromatography (SEC) on a Superdex 200 10/300 GL column (GE Healthcare, Little Chalfont, UK) equilibrated in 50 mM HEPES-NaOH (pH 7.0), 100 mM NaCl, and eluted at 0.5 mL/min with UV detection at 280 nm. Fractions containing Prx2 were pooled, concentrated with Amicon Ultra 10 kDa MWCO, buffer-exchanged, and quantified using a Bio-Rad Protein Assay Kit (Bio-Rad Laboratories, CA, USA) before storage at -80 °C.

*Prx2 oligomer structure and dynamics.*

For both Prx2WT and MT, only oligomeric fractions formed in the presence of PS were collected and concentrated. To investigate the influence of $Mg^{2+}$ and ADP on higher-order assemblies, purified oligomers (20 nM) were incubated at 37 °C for 12 h under three defined conditions: (i) in the presence or absence of 5 mM $MgCl_2$, (ii) with or without 5 mM ADP, and (iii) with a mixture of 5 mM $MgCl_2$, 4.5 mM ADP and 0.5 mM MantADP. Following incubation, samples were loaded onto the SEC column for analysis under the same chromatographic conditions described above.

*Chromatographic fluorescence assay for nucleotide binding.*

An ADP mixture containing 5 mM $MgCl_2$, 4.5 mM ADP, and 0.5 mM Mant-ADP was prepared in 50 mM HEPES-NaOH (pH 7.0) buffer. Mant-ADP (Sigma-Aldrich, St. Louis, MO, USA) was included to enable fluorescence detection. The ADP mixture was incubated separately with Prx2 (0.4 mg/mL) or dioleoyl phosphatidylserine (PS) (0.8 mg/mL; Yuka Sangyo Co., Ltd., Tokyo, Japan) at 37 °C for 12 h. A control reaction containing both Prx2 and PS with the same ADP mixture was prepared and incubated under identical conditions. Following incubation, ADP association was analyzed by SEC using the same Superdex 200 column with in-line UV and fluorescence detection (JASCO system; $\lambda\_exc$ = 355 nm, $\lambda\_em$ = 448 nm).

*Time-course oligomerization and chaperone activity assays*

To monitor oligomerization changes over time, 0.4 mg Prx2 was mixed with 0.8 mg PS in 50 mM HEPES-NaOH pH 7.0. The mixture was aliquoted into separate tubes corresponding to time points from 0 to 4 h and incubated at 37°C. At each time point, samples were subjected to SEC analysis on a same Superdex column, with a flow rate of 0.5 mL/min. Elution profiles were monitored at 280 nm. The retention time window corresponding to oligomeric species (11-13 ml) was recorded. The relative area under the curve was calculated to compare oligomer content across time points. The chaperone activity of oligomer WT and MT were measured using citrate synthase (CS) thermal aggregation assay as described previously [R1].

*HS-AFM analysis*

HS-AFM imaging was performed in aqueous buffer at room temperature using a laboratory-built setup [R2,R3]. A glass stage (1.5 mm x 2 mm) was attached to the Z-scanner, and a mica disc (1.5 mm x 0.05 mm) was glued on top. Imaging was conducted in tapping mode using small cantilevers (BLAC10DS-A2, Olympus, Japan) with a resonant frequency of ~ 0.5 MHz in water, a quality factor of ~ 1.5, and a spring constant of ~ 0.1 N/m. The imaging rate, scan size and feedback parameters were optimized to enable visualization with a minimum tip force. For oligomer imaging, 2 µL of 10 nM Prx2 oligomer was deposited onto freshly cleaved mica, incubated for 10 min, and washed twice with 50 mM HEPES-NaOH (pH 7.0) (imaging buffer) to remove unbound protein before observation. Representative frames are shown in Fig. 3, and complete sequences are provided in Supplementary Movies. For HMW complex observation, we used a carbon-coated mica surface and AFM data were analyzed using custom software, including Kodec, and Umex Viewer, as reported earlier. Oligomer geometry was quantified by calculating the triangular area defined by the centers of the three dimeric units within each hexamer, with vertices corresponding to the highest points relative to the substrate, marking the vertex of each foliole in the HS-AFM imaging. [R2, R3]

R1. Haruyama, T., et al., Negatively Charged Lipids Are Essential for Functional and Structural Switch of Human 2-Cys Peroxiredoxin II. J Mol Biol, 2018. 430(5): p. 602-610.

R2. Uchihashi, T.; Kodera, N.; Ando, T., Guide to video recording of structure dynamics and dynamic processes of proteins by high-speed atomic force microscopy. *Nat Protoc* **2012,** *7* (6), 1193-206.

R3. Ando, T.; Uchihashi, T.; Scheuring, S., Filming biomolecular processes by high-speed atomic force microscopy. *Chem Rev* **2014,** *114* (6), 3120-88.

```
Prx2WT    MASGNARIGKPAPDFKATAVVDGAFKEVKLSDYKGKYVVLFFYPLDFTFVCPTEIIAFSN 60
Prx2MT    MASGNARIGKPAPDFKATAVVDGAFKEVKLSDYKGKYVVLFFYALDFTFVCPTEIIAFSN 60
          ******************************************* ****************

Prx2WT    RAEDFRKLGCEVLGVSVDSQFTHLAWINTPRKEGGLGPLNIPLLADVTRRLSEDYGVLKT 120
Prx2MT    RAEDFRKLGCEVLGVAVDSQFTHLAWINTPRKEGGLGPLNIPLLADVTRRLSEDYGVLKT 120
          ***************:********************************************

Prx2WT    DEGIAYRGLFIIDGKGVLRQITVNDLPVGRSVDEALRLVQAFQYTDEHGEVCPAGWKPGS 180
Prx2MT    DEGIAYRGLFIIDGKGVLRQITVNDLPVGRSVDEALRLVQAFQYTDEHGEVCPAGWKPGS 180
          ************************************************************

Prx2WT    DTIKPNVDDSKEYFSKHN  198
Prx2MT    DTIKPNVDDSKEYFSKHN  198
          ******************
```

**Fig. S1. Sequence alignment between Prx2 WT and Prx2 MT.**
Two-point mutations (Pro44→Ala, Ser76→Ala) are highlighted in red.

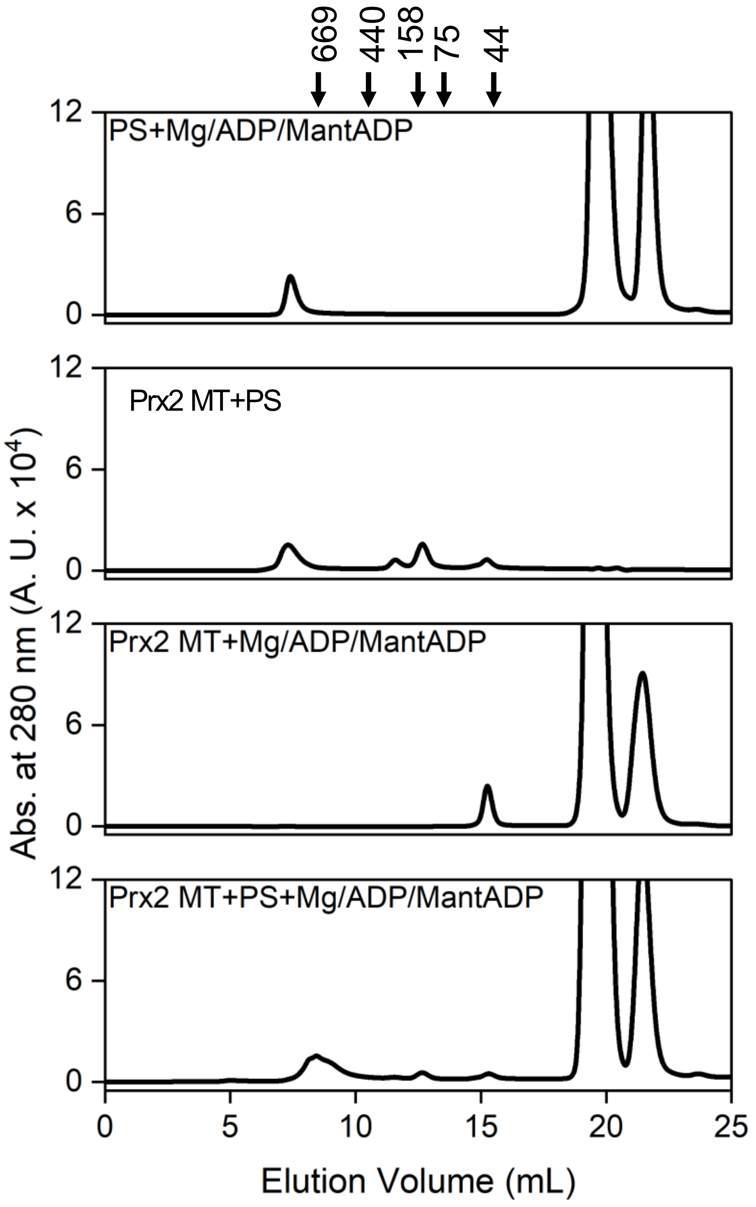


**Fig. S2**. **SEC chromatograms of the Prx2$_{P44A/S76A}$ mutant under different conditions.** SEC chromatograms of the Prx2 P44A/S76A mutant under the indicated conditions. An early-eluting HMW fraction was observed when MT Prx2, PS, $Mg^{2+}$, ADP, and Mant-ADP were combined.

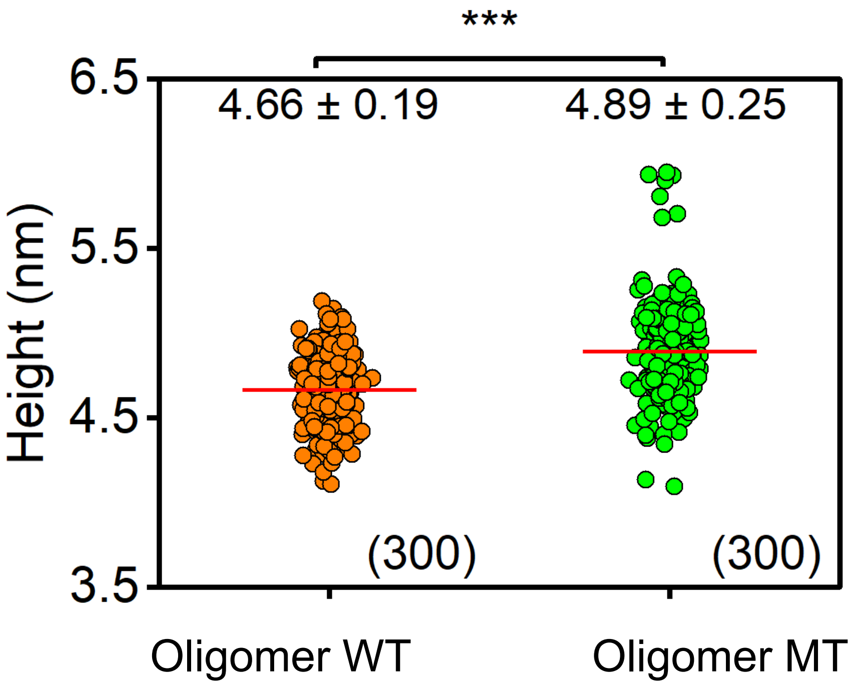


**Fig. S3. Height analysis of Prx2 oligomers.**

Scatter plot showing the height distribution of Prx2 WT (orange, n=300) and Prx2 MT (green, n=300) oligomers analyzed from HS-AFM images. Red lines indicate mean values. (Oligomer WT = 4.66 ± 0.19 nm, Oligomer MT = 4.89 ± 0.25 nm for mean height ± SD). Statistical analysis by Student's t-test confirmed a significant difference between WT and MT (*** $p < 0.001$).

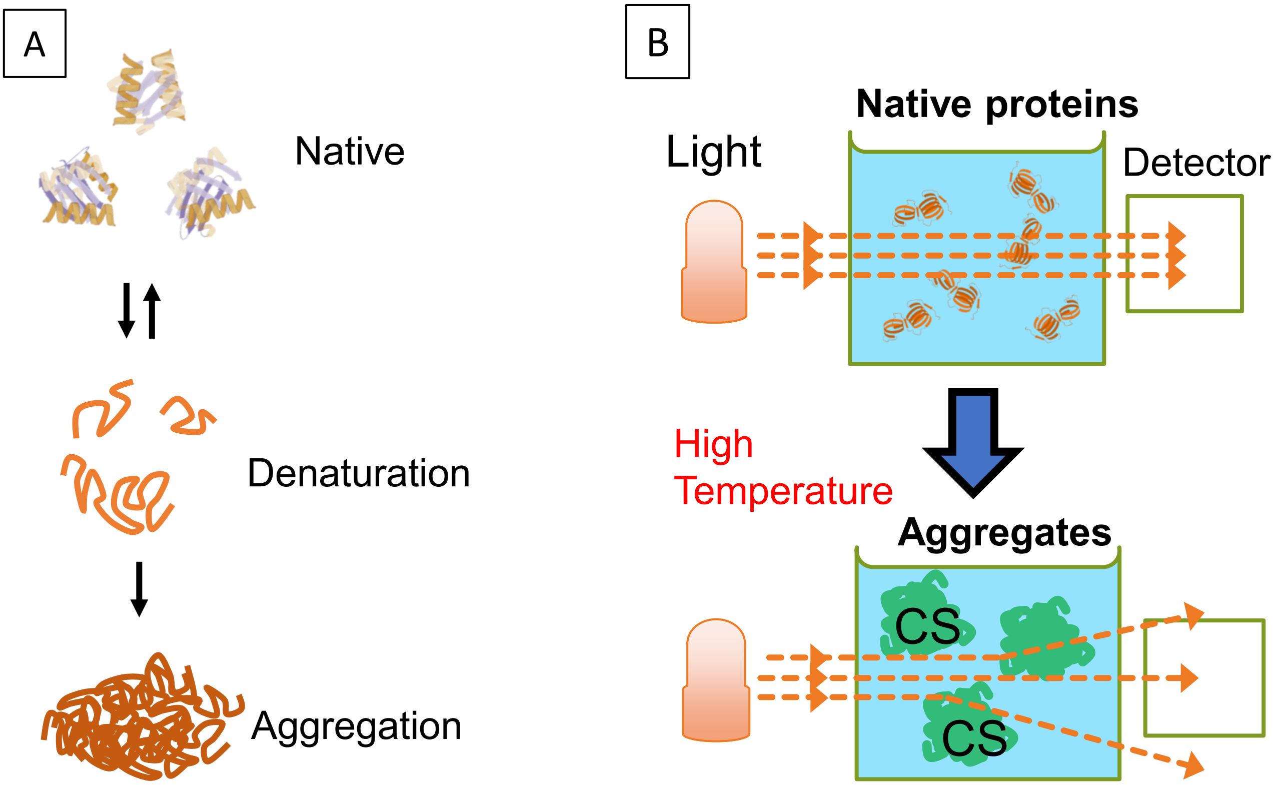


**Fig. S4. Scheme illustrating the principle of the citrate synthase aggregation assay.**
(A) Schematic diagram of protein structural states. The top, middle, and bottom illustrations represent the native, denatured, and aggregated states of the protein, respectively. (B) Native CS maintains a compact folded structure with minimal light scattering (top). Under high temperature, CS proteins undergo denaturation and aggregation, forming large complexes that scatter light strongly (bottom). Chaperone activity of Prx2 is determined through its ability to prevent CS aggregation at high temperatures.

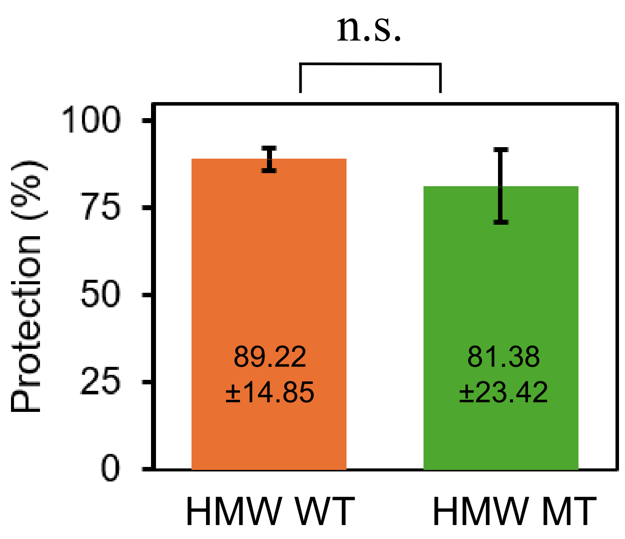


**Fig. S5. Chaperone activities of WT and MT Prx2 HMW assemblies.**
The protective capacities of WT (orange) and MT (green) HMW assemblies were 89.22±14.85% and 81.38±23.42%, respectively. No statistically significant difference was observed between the two groups (n.s.).

**Results:**

- RMSD = 0.07 Å

- P44 Cα displacement = 0.136 Å
- S76 Cα displacement = 0.122 Å

- P44 angle:

WT = 104.35°
MT = 104.58°
Δ = 0.23°

- S76 angle:

WT = 130.77°
MT = 132.56°
Δ = 1.79°

- WT H-bond distance:

TYR43 OH = 3.09 Å
ASP46 OD1 = 3.22 Å
ASP78 OD2 = 3.29 Å
HIS83 NE2 = 3.48 Å

**Atom around OG of S76:**

A 43 TYR CE2
A 43 TYR CZ
A 43 TYR OH
A 46 ASP OD1
A 76 SER N
A 76 SER CA
A 76 SER C
A 76 SER O
A 76 SER CB
A 76 SER OG
A 77 VAL N
A 78 ASP N
A 78 ASP CB
A 78 ASP OD2
A 83 HIS CE1
A 83 HIS NE2

**Table 1. Data sets of structural comparison of AlphaFold-predicted WT and P44A/S76A hPrx2.**
AlphaFold-predicted WT and MT structures were superimposed, yielding an RMSD of 0.07 Å, indicating preservation of the global fold. The Cα displacements at positions P44 and S76 were 0.136 Å and 0.122 Å, respectively. Local backbone geometry was minimally affected, with bending angle changes of 0.23° around P44 and 1.79° around S76. Analysis of the WT model suggested that S76 participates in a local hydrogen-bonding network.

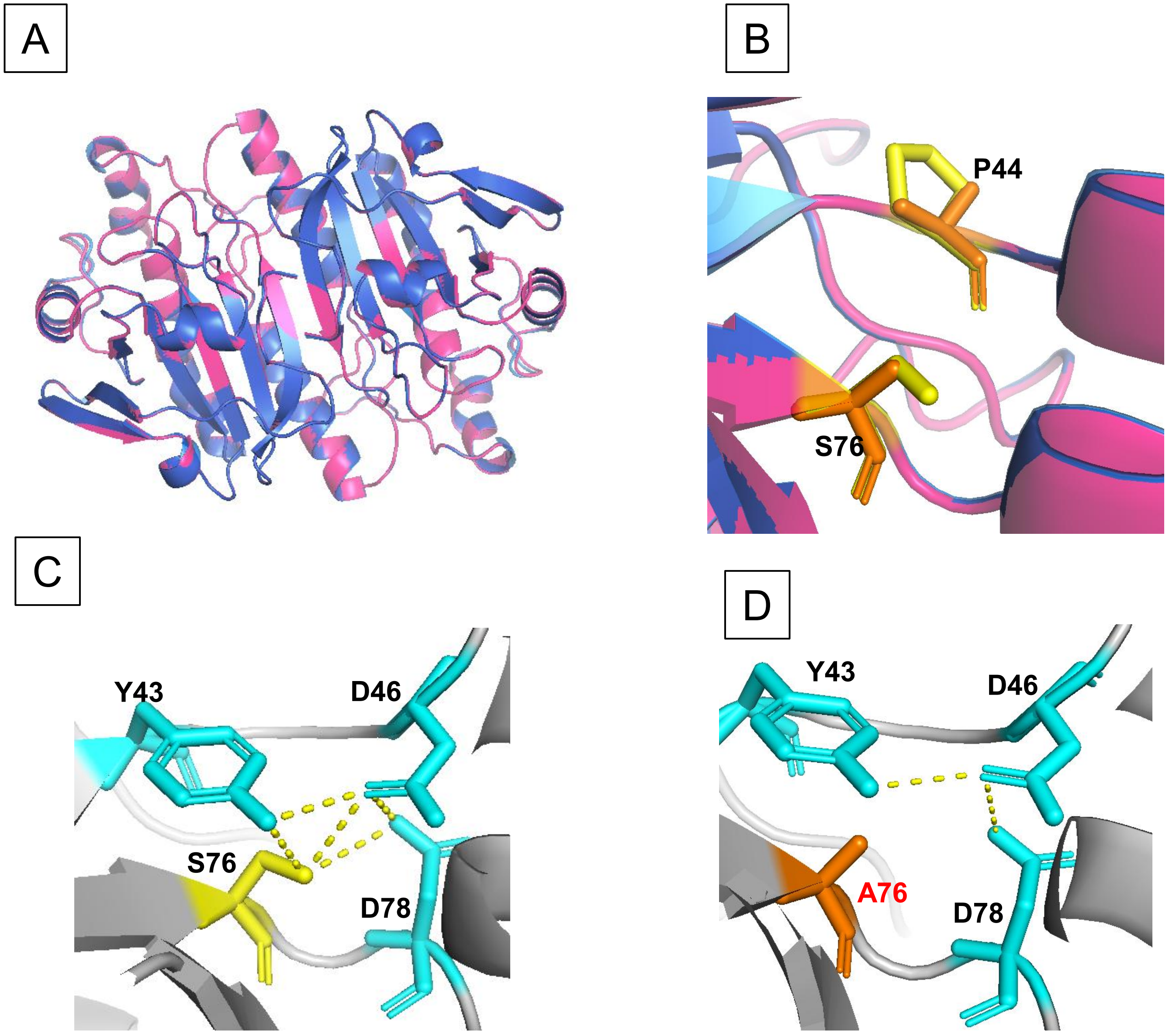


**Figure S6. Structural comparison of AlphaFold-predicted WT and P44A/S76A hPrx2.**
(A) Structural superposition of WT (blue) and MT (magenta) models showing a backbone RMSD of 0.07 Å. (B) Local view of residues P44 and S76 with WT (yellow) and MT (orange).
(C) Predicted hydrogen-bonding network surrounding S76 in the WT structure. Potential contacts were observed between S76 and Y43 (3.09 Å), D46 (3.22 Å), D78 (3.29 Å), respectively. Besides, the interaction between D46 and Y43 was 2.96 Å, D46 and D78 was 3.15 Å. (D) Substitution to A76 removes the side-chain hydroxyl group, thereby disrupting S76-mediated contacts while preserving the overall local structural organization.

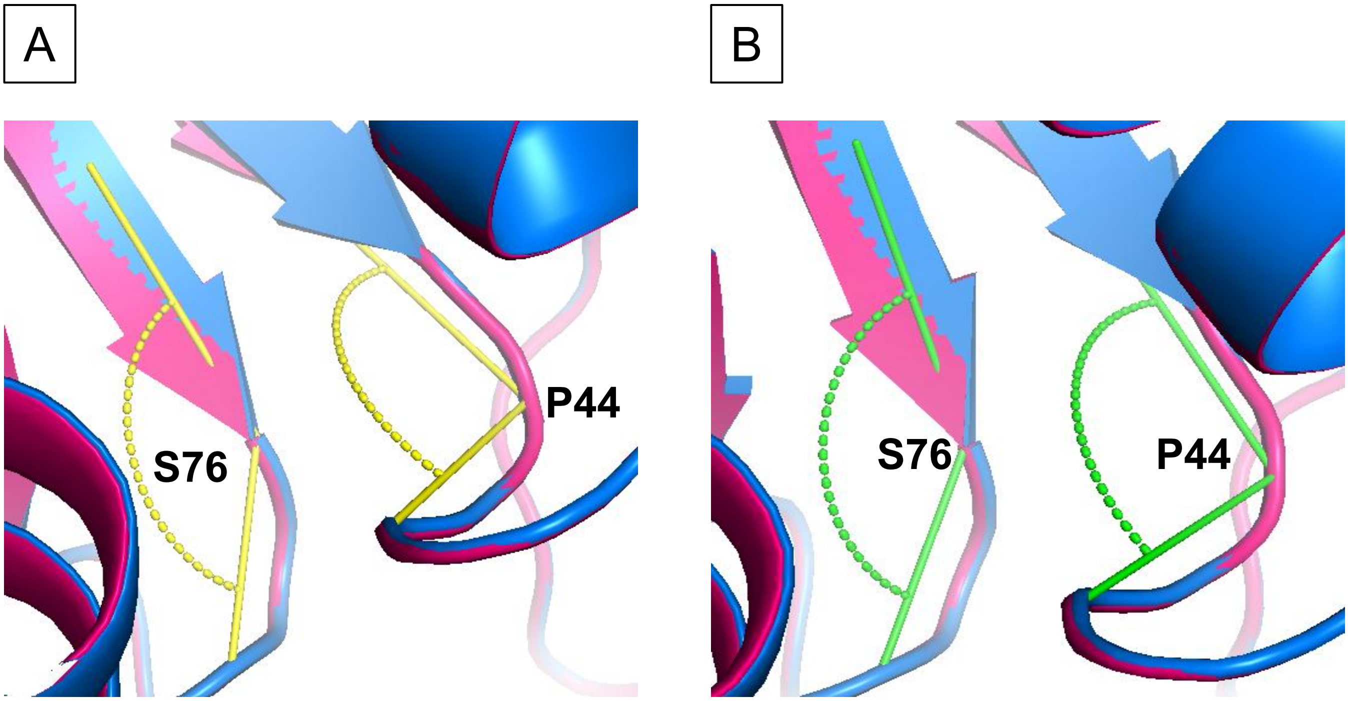


**Figure S7. Backbone bending angle comparison of (A) Prx2 WT and (B) Prx2 MT.** The backbone bending angles changed from 104.35$^{o}$ (yellow) to 104.58$^{o}$ (green) around P44 and from 130.77$^{o}$ (yellow) to 132.56$^{o}$ (green) around S76.

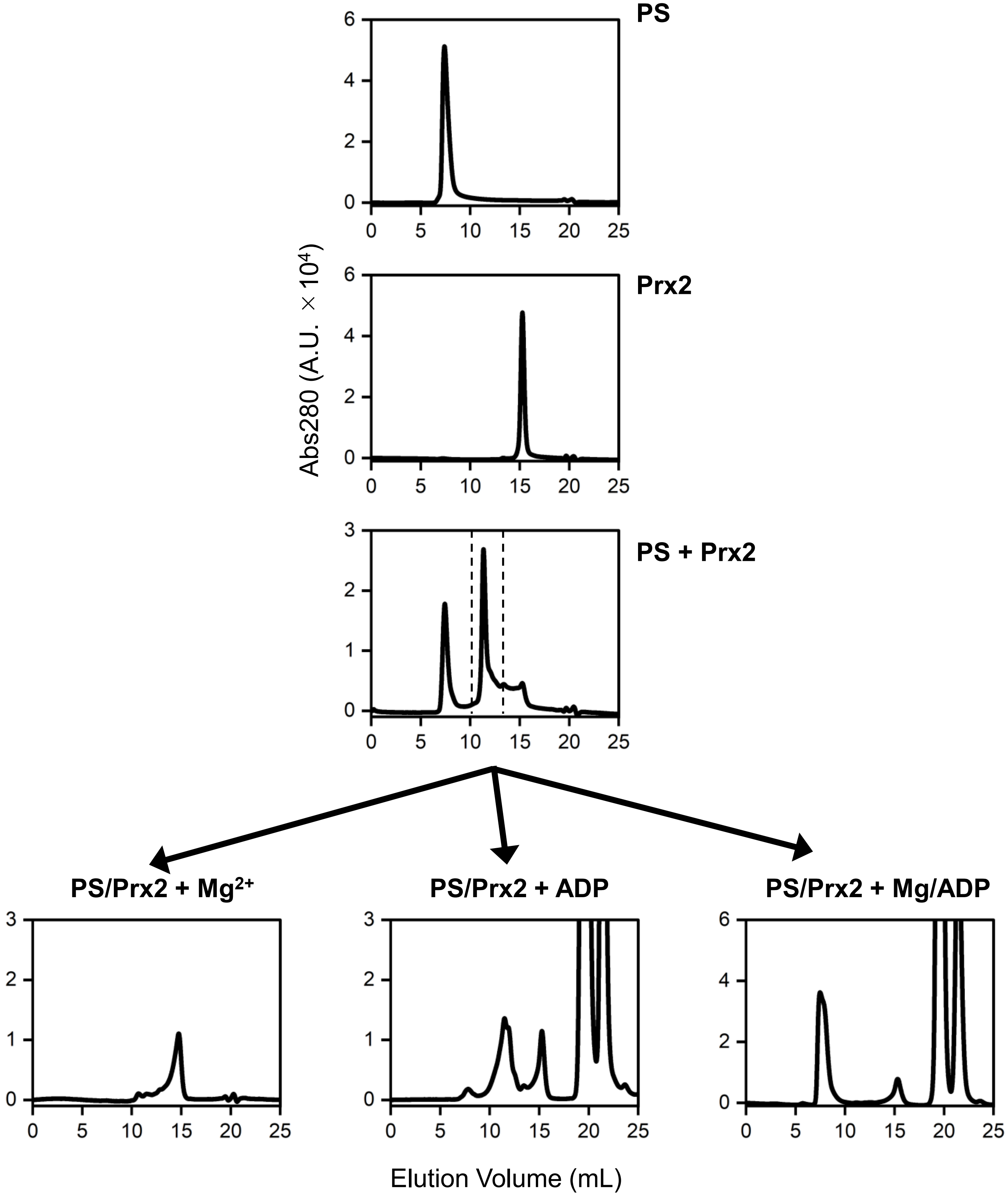


**Fig. S8. Experimental procedure for confirming ADP and $Mg^{2+}$ involvement in HMW complex formation.** The peak indicated by the dotted line (Prx2-PS complex) was collected and fractioned.

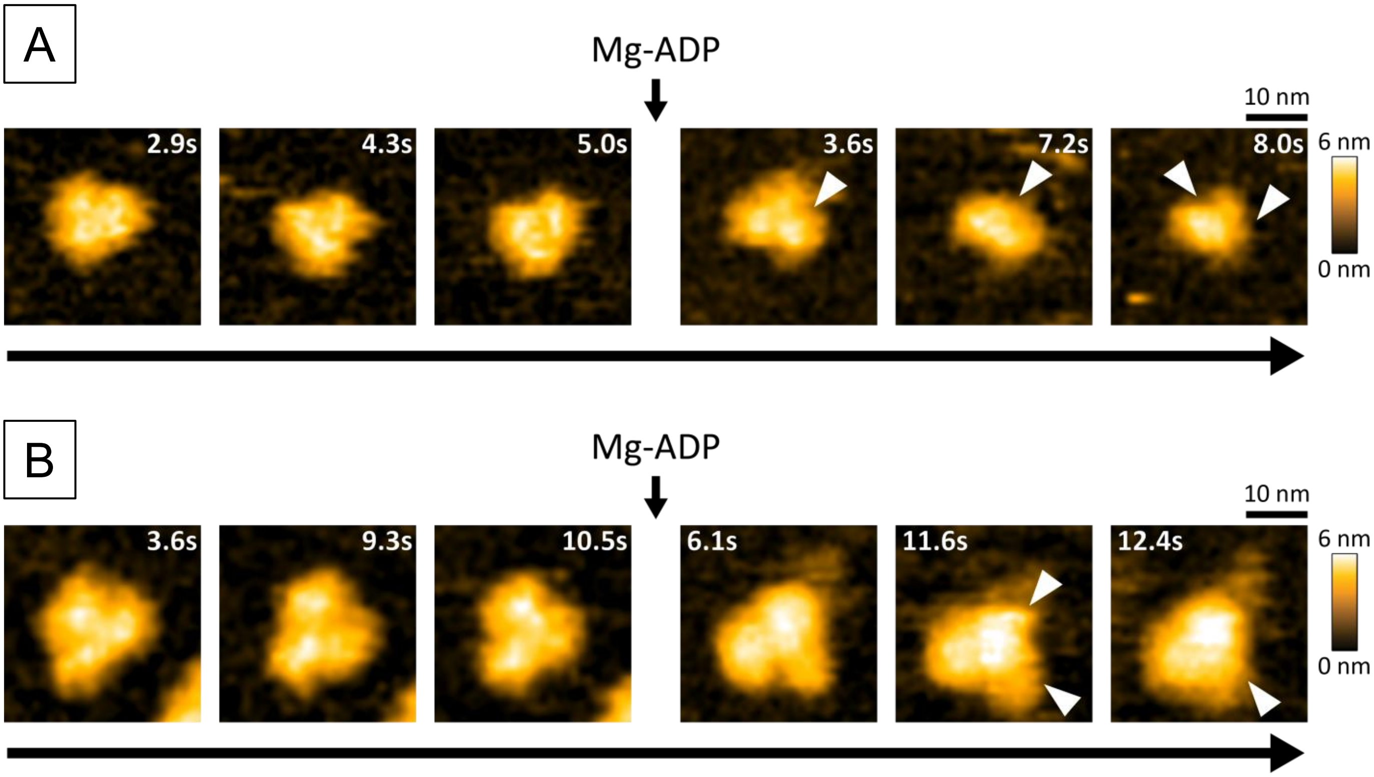


**Fig. S9 HS-AFM time-lapse imaging of oligomers (A) WT and (B) MT before and after adding Mg-ADP.** A) WT and (B) MT oligomers. Mg-ADP and imaging buffer were preincubated at 37 ° C before addition to the imaging chamber. Time stamps before Mg-ADP addition indicate elapsed imaging time, whereas those after addition are referenced to the time of Mg-ADP addition (t = 0). White arrowheads indicate regions of oligomer fragmentation. Scale bar: 10 nm; Z-scale: 6 nm; scan size: 50 × 50 nm² (50 × 50 pixels).

**Supplementary movie 1**. **Time-lapse HS-AFM movie showing structural dynamic of oligomer WT.** Z-scale was 6 nm. Scanning area was 50 x 50 $nm^2$ with 50 x 50 pixels. Imaging rate was 0.1 s/frame (~10 fps).

**Supplementary movie 2**. **Time-lapse HS-AFM movie showing structural dynamic of oligomer MT.** Z-scale was 6 nm. Scanning area was 50 x 50 $nm^2$ with 50 x 50 pixels. Imaging rate was 0.1 s/frame (~10 fps).

**Supplementary movie 3**. **Time-lapse HS-AFM movie showing the degradation of WT oligomers in the presence of $MgCl_2$**. Z-scale was 6 nm. Scanning area was 50 x 50 $nm^2$ with 50 x 50 pixels. Imaging rate was 0.15 s/frame (~7 fps).

**Supplementary movie 4**. **Time-lapse HS-AFM movie showing the degradation of MT oligomers in the presence of $MgCl_2$**. Z-scale was 6 nm. Scanning area was 50 x 50 $nm^2$ with 50 x 50 pixels. Imaging rate was 0.1 s/frame (~10 fps).

**Supplementary movie 5. Time-lapse HS-AFM movie showing the degradation of WT oligomers in the presence of ADP.** Z-scale was 6 nm. Scanning area was 50 x 50 $nm^2$ with 50 x 50 pixels. Imaging rate was 0.1 s/frame (~10 fps).

**Supplementary movie 6**. **Time-lapse HS-AFM movie of the interaction between MT oligomers and ADP.** Z-scale was 6 nm. Scanning area was 50 x 50 $nm^2$ with 50 x 50 pixels. Imaging rate was 0.1 s/frame (~10 fps).

**Supplementary movie 7. Time-lapse HS-AFM movie showing the degradation of WT oligomers in the presence of Mg-ADP.** Z-scale was 6 nm. Scanning area was 50 x 50 $nm^2$ with 50 x 50 pixels. Imaging rate was 0.2 s/frame (~5 fps).

**Supplementary movie 8. Time-lapse HS-AFM movie showing the degradation of MT oligomers in the presence of Mg-ADP.** Z-scale was 6 nm. Scanning area was 50 x 50 $nm^2$ with 50 x 50 pixels. Imaging rate was 0.1 s/frame (~10 fps).
